# G4All: a database of experimentally confirmed G-quadruplex-forming sequences

**DOI:** 10.64898/2026.08.05.743020

**Authors:** Parth Rajput, Anne Cucchiarini, Dmitrii Trubetskoy, Giacomo Ferrari, Yun Chen, Muafia Baigum, Lionel Guittat, Laurent Lacroix, Jean-Louis Mergny

**Affiliations:** Laboratoire d’Optique et Biosciences, École Polytechnique, CNRS, Inserm, Institut Polytechnique de Paris, 91120 Palaiseau, France; Indian Institute of Technology Delhi, New Delhi, India; Department of Biochemistry, University of Agriculture Faisalabad, Faisalabad 38040, Pakistan; Université Sorbonne Paris Nord, UFR SMBH, 93000 Bobigny, France; Institut de Biologie de L’École Normale Supérieure (IBENS), École Normale Supérieure, CNRS, INSERM, Université PSL, Paris, France

**Keywords:** G-Quartet, Database, Prediction, Melting temperature, Interactive Website, G4Hunter, Aptamer

## Abstract

G-quadruplexes (G4s) are non-canonical nucleic acid structures with critical roles in gene regulation, genomic stability, and disease, making them prime targets for therapeutic and biotechnological applications. However, the absence of a curated, centralized, experimentally validated repository of short (mostly synthetic) G4-forming sequences, paired with appropriate single-stranded or hairpin controls, has limited reproducibility and hindered progress in the field. Here, we introduce G4All, a comprehensive, curated database of G4-forming short DNA and RNA sequences, complemented by rigorously selected non-G4 controls, all studied under roughly similar experimental conditions (around 100 mM potassium ion at near neutral pH). G4All consolidates sequences validated by diverse experimental methods, including circular dichroism, NMR, UV spectroscopy, with standardized annotations for topology, thermal stability, and experimental conditions. By providing both positive and negative datasets, G4All enables rigorous comparative analyses, assay benchmarking, and the development of predictive models. The database supports a broad range of applications, from fundamental studies of G4 biophysics to the rational design of aptamers, as well as benchmarking prediction algorithms. Future developments will expand G4All to include user-submitted datasets and more RNA sequences. Freely accessible, G4All offers a searchable interface and downloadable datasets, establishing a community-driven hub to accelerate discovery and standardization in G4 research.

**GRAPHICAL ABSTRACT:** 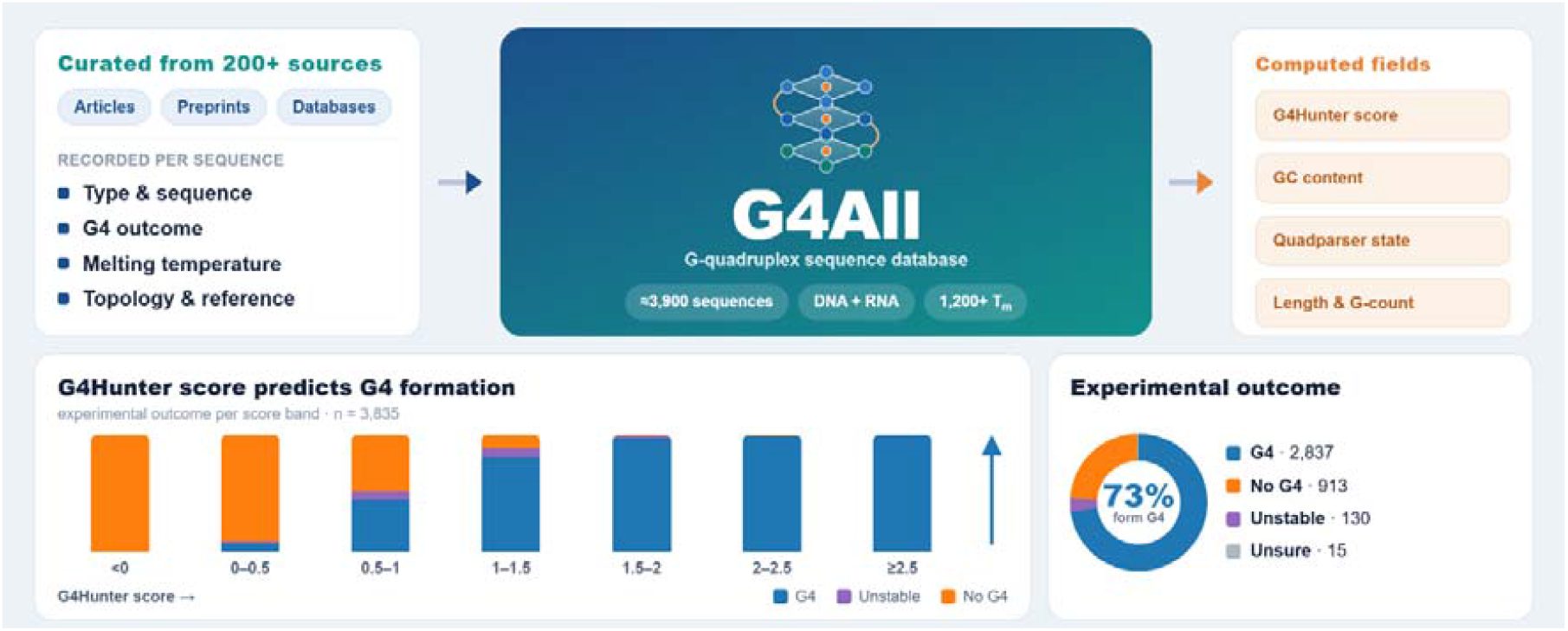

## INTRODUCTION

G-quadruplexes (G4s) are non-canonical nucleic acid secondary structures formed by guanine-rich DNA and RNA sequences that fold into stable four-stranded helices stabilized by Hoogsteen hydrogen bonds and cations such as potassium (for a review : 1). Over the past two decades, compelling evidence has demonstrated that G4s play pivotal roles in key biological processes, including gene regulation, telomere maintenance, DNA replication, and genomic stability, as well as in pathological conditions such as cancer and neurodegenerative diseases (for a review: 2). Their prevalence in functionally critical genomic regions—such as promoters, telomeres, origins of replication, and untranslated regions—highlights their potential as therapeutic targets and regulatory elements. Furthermore, the dynamic formation and resolution of G4s in both DNA and RNA underscore their significance in transcriptional and post-transcriptional regulation, as well as in the maintenance of genomic integrity.

Despite their biological and biomedical relevance, the systematic study of G4s has been hindered by the lack of comprehensive, curated, and experimentally validated datasets. While computational predictions have identified millions of putative G4-forming sequences (PQS) across genomes, experimental validation remains sparse and scattered across disparate studies, often lacking standardized controls or consistent methodological frameworks. This gap limits the reproducibility of research, impedes the development of G4-targeting tools, and complicates the interpretation of high-throughput sequencing data.

To address this challenge, we present **G4All**, a specialized database dedicated to G4-forming short DNA and RNA sequences, complemented by a curated collection of non G4-forming controls (single-strands, hairpins, i-motifs). Some of these sequences may be natural, found in a variety of eukaryotic, prokaryotic or viral genomes; others may be model sequences used to investigate sequence effects on G4 structure and stability. The whole database can be downloaded as a single file at https://doi.org/10.5281/zenodo.21216943, and is freely interrogable at https://g4-all.streamlit.app. it consolidates experimentally verified G4-prone sequences alongside non-G4-forming counterparts, providing researchers with a robust resource for comparative analyses, method benchmarking, and hypothesis-driven investigations. By integrating data from diverse experimental techniques, such as circular dichroism, UV spectroscopy (TDS, IDS), or G4 specific fluorescent probes, G4All offers a standardized and searchable repository to accelerate discovery in the G4 field. This resource not only facilitates the design of targeted experiments but also supports the development of computational models for G4 prediction and the exploration of G4s as therapeutic targets. When available, information on thermal stability (T_*m*_) and topology in potassium is also provided. Here, we describe the scope, architecture, and utility of G4All, emphasizing its role in advancing the study of these fascinating nucleic acid structures.

## MATERIAL AND METHODS

### Building the database

Figure 1 summarizes the processes used to build G4All. Step 1 and 2 involved a repeated process of collecting research papers, preprints, or unpublished results from our group which analyzed short G4 and non G4 forming sequences *in vitro* and then extracted and compiled data from these sources. Step 3 and 4 subsequently involve cleaning the collected data, adding new calculated fields and analysis of relations between different fields. To find other G4—related databases to which G4All was compared were identified via a literature search on Pubmed, and by queries to Biomni (3).

**Figure 1:**
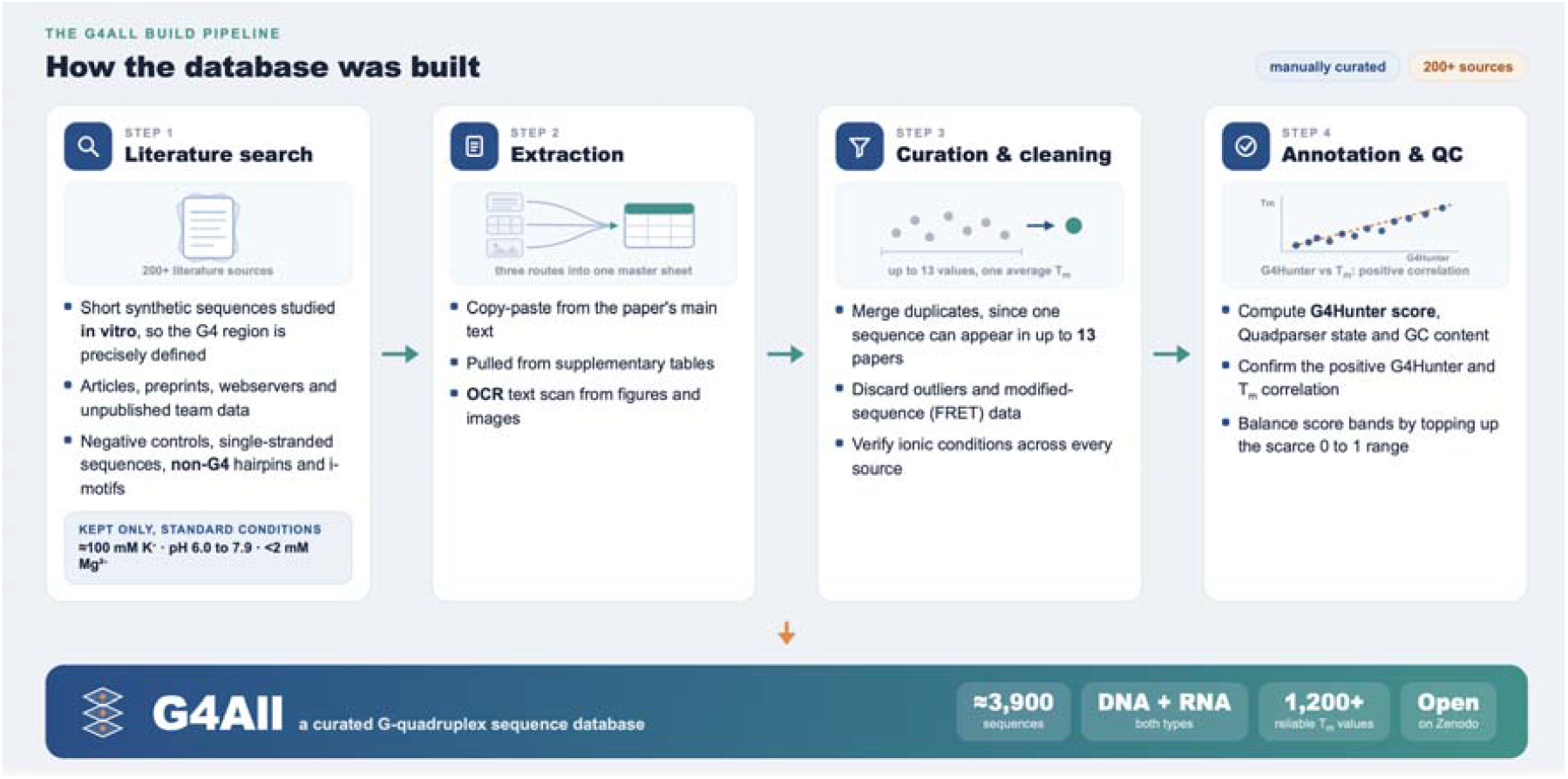
Building G4All. The analysis in step 4 helped to find biases in existing literature.

### Criteria for selection or inclusion

The database is composed of relatively short synthetic sequences that were analyzed *in vitro*. A significant part of the data (over 2200 sequences) was collected by us over the years, starting 30 years ago, when we found out that recording absorbance changes at 295 nm allowed to follow G4 thermal denaturation (4). Yet, over 270 different sources have been used so far to build this repository – the origin of the data provided for all sequences is provided online. We did not consider motifs purely identified *in silico*, or motifs embedded into long genomics / transcriptomics data (as in G4Access or BG4-ChiP Seq experiments). Focusing on short, synthetic sequences characterized *in vitro* offers distinct advantages over genomic or transcriptomic data. *In vitro* studies allow precise control of experimental conditions (*e*.*g*., ionic strength, pH, temperature), eliminating the confounding variables of cellular environments, such as molecular crowding, protein interactions, or supercoiling, that obscure fundamental biophysical properties. It also excludes possible competition with hairpin formation with complementary sequences nearby that can be addressed separately. Our initial goal is to study the G4 itself, and integration of longer motifs may be considered later. Short sequences provide standardized, reproducible models for benchmarking assays, validating computational predictions, and designing G4-targeting tools, while their simplicity enables mechanistic insights into topology, stability, and ligand interactions that are difficult to extract from complex, context-dependent genomic data. This reductionist approach thus establishes a reliable foundation for developing practical applications.

For the sequences reported to form G4s (stable or not), we kept only the motifs for which intramolecular formation is possible, at least in theory. This excludes very short motifs such as GGGTTTGGG which form intermolecular G4s and sequences with less than 8 guanines, therefore unable to form 2 quartets in an intramolecular fashion. Of note, we took great care in keeping data obtained under standard conditions (≈ 100 mM K^+^ at near neutral pH; available for over 1200 sequences), to allow meaningful comparisons, especially when considering T_*m*_ values Among sequences able to form G-quadruplex structures, we made a distinction between those thermally unstable (“Unstable G4s” for which T_*m*_ < 37°C) for which the oligonucleotide will be predominantly unfolded at human physiological temperature, and other G4-forming motifs (called “G4s”). Note that some articles reported G4s as “unstable”, even if their T_*m*_ was reported to be above 37°C: these quadruplexes were relabeled as “G4s”.

Negative controls are composed of sequences for which experimental evidence exists that the oligonucleotide remains single-stranded, or adopts a different structure, such as a hairpin, intramolecular triplex, or i-DNA. Some of these controls contain few or no guanines, others may be relatively GC-rich and prone to Watson-Crick pairing. A number of hairpins were included: these sequences have a G4Hunter score close to zero. The most reliable methods to conclude on G4 formation *in vitro* are to compare melting/IDS/TDS/CD profiles in the presence or absence of a G4-promoting ion such as a potassium.

### Curating data

Sequence curation included removing space, dots or hyphens between nucleotides. Degenerate nucleotides such as H or N were kept. Most sequences incorporating modifications were removed: the database is composed mostly of unmodified DNA or RNA sequences that did not contain a high number of non-natural nucleotides. In most cases in which a G4 is formed, molecularity is not established. As discussed above, we aimed to build a database of intramolecular G4s and we therefore discarded sequences for which this type of folding is not possible (*i*.*e*., those which can only form intermolecular G4). Finally, we sometimes had to determine the reverse complement of a given sequence when G4 formation was reported for a C-rich motif.

Of note, when considering intramolecular structures, T_*m*_ and stability are independent of strand concentration, meaning that the results obtained at different DNA concentrations can still be compared. We then checked the experimental conditions used, in order to gather data obtained under relatively similar, if not identical conditions. Our reference conditions are 100 mM potassium at near-neutral pH. For this reason, we considered results obtained in 80-120 mM K^+^ in the pH range 6.0 to 7.9 (contrary to i-DNA, G4 formation is relatively insensitive to pH in that range). A number of sequences have been studied by different groups. When T_*m*_ values were reported, we generally keep the average, unless outlier values were identified; manual checking of each experiment is then necessary.

### Parameters listed

The following information is provided in the G4All database:

- Type (DNA/RNA),
- Sequence (5’-3’),
- G4Hunter score (between -4 and 4),
- G4Hunter (G4Hmax) “*max score*” (for long sequences, it corresponds to the highest score for any window of 25 nt within the motif. For example, a core G-quadruplex with a 5’ or 3’ single-stranded extension such as T_30_(G_3_TTA)_3_G_3_ has a G4H score of 0.71, but a G4H”max” score of 1.44).
- Conclusion: Evidence for G4 formation (G4 / No G4 / unstable G4 when T_*m*_ <37°C / or “unsure” for a few sequences for which contradictory results were reported). Approximately 76% of the sequences were found to form quadruplexes, stable or not.
- “Final” T_*m*_ value in standard conditions, sometimes averaged from different sources (up to 17 independent determinations!). Reliable values were collected for over 1,200 G4 motifs. An arbitrary T_m_ value of 95°C was assigned to all sequences for which T_m_ was reported as above 90°C or 95°C.
- Number of independent T_m_ values used to calculate this average (after outlier removal).
- Length (in nucleotides),
- Quadparser (QP) compliance (Yes/No) indicates whether the sequence would be picked by QP with runs of at least 3Gs, and loops of 1-7 nucleotides.
- GC-content (in %)
- PDB code, for the intramolecular G4 structures that were determined in potassium conditions
- Total G count
- Notes / additional comments
- Topology, as determined by CD or NMR / Crystallography. It includes “r” values as used by some papers
- Name in the corresponding paper, if provided.
- Source (in the paper) – for example “Table 1”
- Study type (reference) and methods used for characterization.
- User (for internal purpose; person who added the sequence in the database)

**Table 1:** G4All and other databases. *Intra- or inter-refers to molecularity. *: assumed, not systematically demonstrated*.

| Name<br>Reference<br>or URL | DNA<br>/<br>RNA | #<br>Entries | Methods | Intra /<br>Inter | Standard<br>conditions | Non G4<br>controls | Focus / Note |
| --- | --- | --- | --- | --- | --- | --- | --- |
| **G4All**<br>g4-all.streamlit.app/<br>*This study* | Both | 3900 | Biophysic<br>s | Intra* | Yes | Yes | G4 *in vitro* by<br>short synthetic<br>sequences under<br>standard<br>conditions |
| **G4LDB**<br>g4ldb.com (6,7) | Both | 4800<br>ligands | Variable | varies | No | No | G4 ligands |
| **G4IPDB**<br>people.iiti.ac.in/~ami<br>tk/bsbe/ipdb (8) | Both | 200<br>proteins | Variable | varies | No | No | G4-binding<br>proteins |
| **OnQuadro**<br>onquadro.cs.put.poz<br>nan.pl (9) | Both | 518<br>G4s | PDB-deposited<br>struct. | varies | No | No | Structural<br>information |
| **DSSR-G4DB**<br>g4.x3dna.org<br>(no reference) | Both | >500 | NMR,<br>Xal | varies | No | No | Structural<br>information<br>(some sequences<br>collected in G4All) |
| **G4Atlas**<br>g4atlas.org<br>(10) | RNA | Many | Transcrip<br>tome-<br>wide<br>rG4s | Intra* | No | No | RNA only |
| **QUADRAtlas**<br>quadratlas.scottgrou<br>p.med.usherbrooke.ca (11) | RNA |  | Experime<br>ntal &<br>Predict. |  | No | No | Includes RNA G4<br>binding proteins |
| **Quadbase2**<br>(12-13) | DNA |  | Predict. | Intra* | No | No | Eukaryotes |
| **EndoQuad**<br>EndoQuad.chenzxlab<br>.cn (14) | DNA |  | Predict. | Intra* | No | No | Genomic<br>annotation |
| **GRSDB2**<br>n/a<br>(15) | RNA | Many | Predict. | Intra* | No | No | pre-mRNAs and<br>mRNAs |
| **Greglist**<br>(16) | DNA | Many | Predict. | Intra* | No | No | Promoters |
| **G4RNA**<br>scottgroup.med.ushe<br>rbrooke.ca/G4RNA<br>(17) | RNA | 334 | Structura<br>l probing<br>&<br>biophysic<br>s | Intra* | No | No | (some sequences<br>collected in G4All) |
| **GAIA**<br>gaia.cobius.usherbro<br>oke.ca (18) | Both | Many | Structura<br>l probing<br>&<br>biophysic<br>s | Intra* | No | No | (some sequences<br>collected in G4All) |
| **G4Bank**<br>tubic.tju.edu.cn/g4b<br>ank<br>(19) | DNA | Many | Genome-wide<br>Incl.<br>Predict. | Intra* | No | No | DNA only |
| **Non-B DB**<br>nonb-<br>abcc.ncifcrf.gov<br>(20) | DNA | Many | Predictio<br>ns | Intra* | No | No | Not only G4s, but<br>Z-DNA,<br>triplexes... |
| Quadrupia<br>(21) | Both | Many | Predictions | Intra* | No | No | Whole genome predictions |

## RESULTS

### Brief description of the current version

As of today, G4All includes ≈ 3,900 DNA and RNA sequences (mean length: 28.0 ± 9.5 nucleotides; max 99; min 10; **Figure 2**) from 272 different sources (articles, reviews, preprints, databases, unpublished data from our group). G4-forming and non G4-forming control sequences had similar lengths (averages = 27 and 30, respectively). All results have been manually and carefully curated (see experimental section).

**Figure 2:**
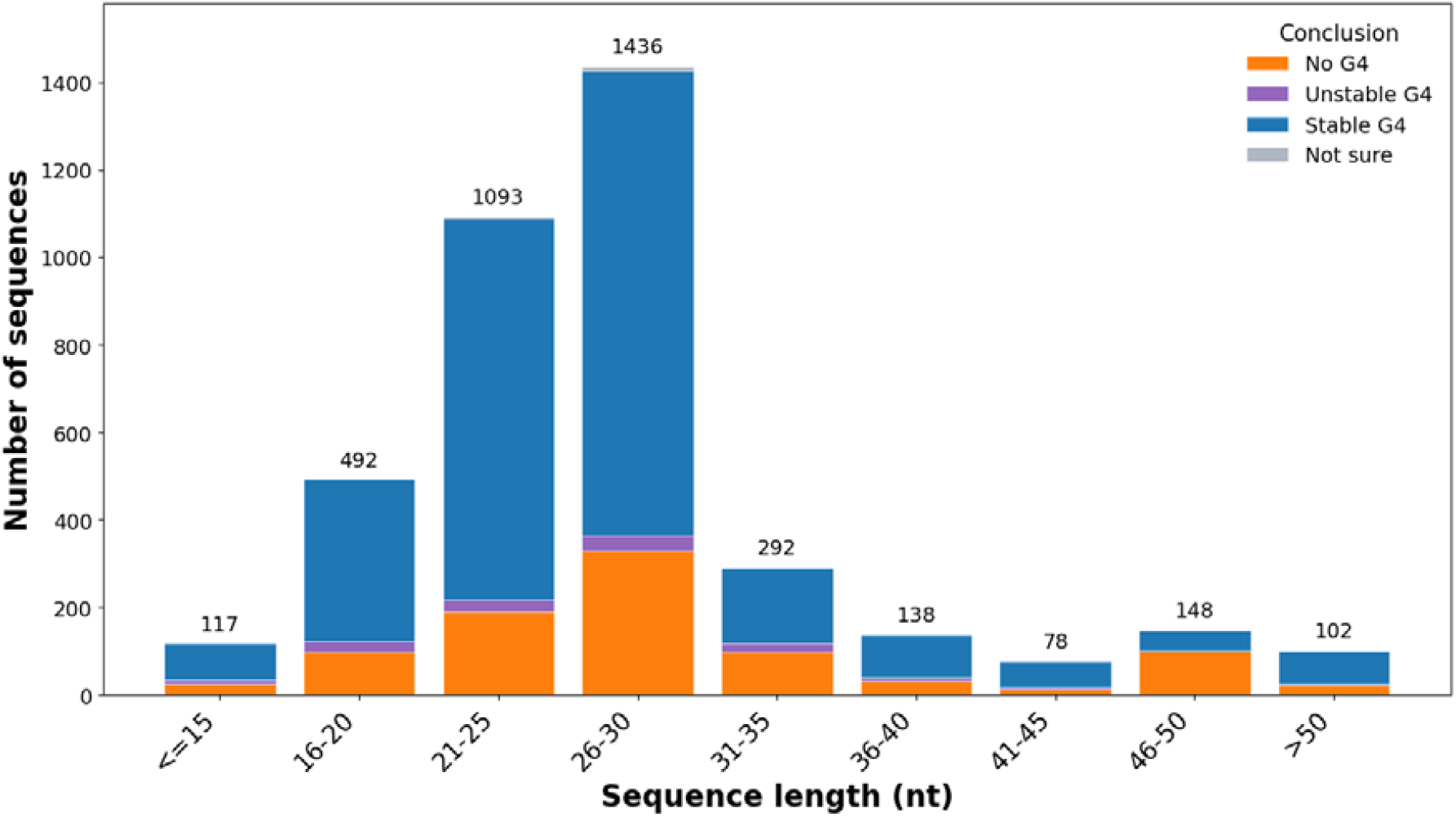
Length of the sequences included in the database. Total number for each size range is provided on top. G4-forming, Unstable G4s (with T_m_ <37°C) and non G4 forming sequences are shown in blue, purple and orange, respectively. A few sequences for which no conclusion could be drawn are shown in grey (*e*.*g*., when two articles report opposite results).

### Accessing, exploring and downloading the database

Figure 3 corresponds to a screenshot of the G4All browser. The user-friendly interface can be used to easily search for particular sequences, threshold ranges, types. The distributions and relationships tabs allow the visualization of the collected G4All dataset by comparing pairwise fields.

**Figure 3:**
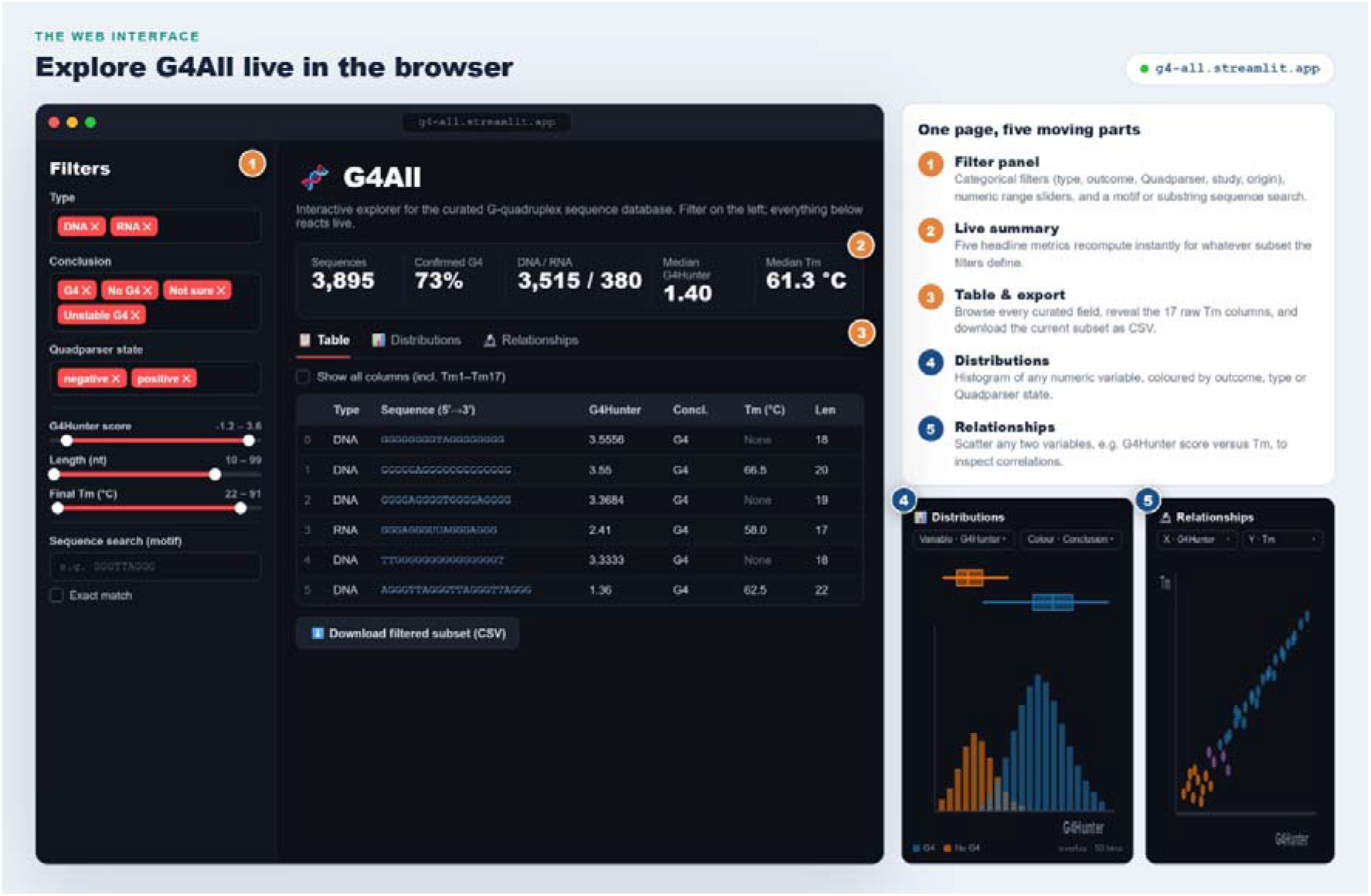
The G4All Web interface. Accessible at: g4-all.streamlit.app

### G4Hunter accuracy

The amount of data collected here allowed us to test the accuracy of G4Hunter on a dataset approximatively 20 times larger than the one used at the time of the original G4Hunter study (5). **Figure 4** illustrates the conclusion reached for all the sequences tested, ranked by G4Hunter score. This data confirms that a threshold of 1.15 to 1.2 optimizes accuracy (close 90%) for DNA. If the goal is to minimize false negatives, higher thresholds may be chosen: 98.6% of all sequences with a score above 1.25 form a quadruplex, stable or not, while 99.6% of all sequences with a score above 1.5 form a stable quadruplex.

**Figure 4:**
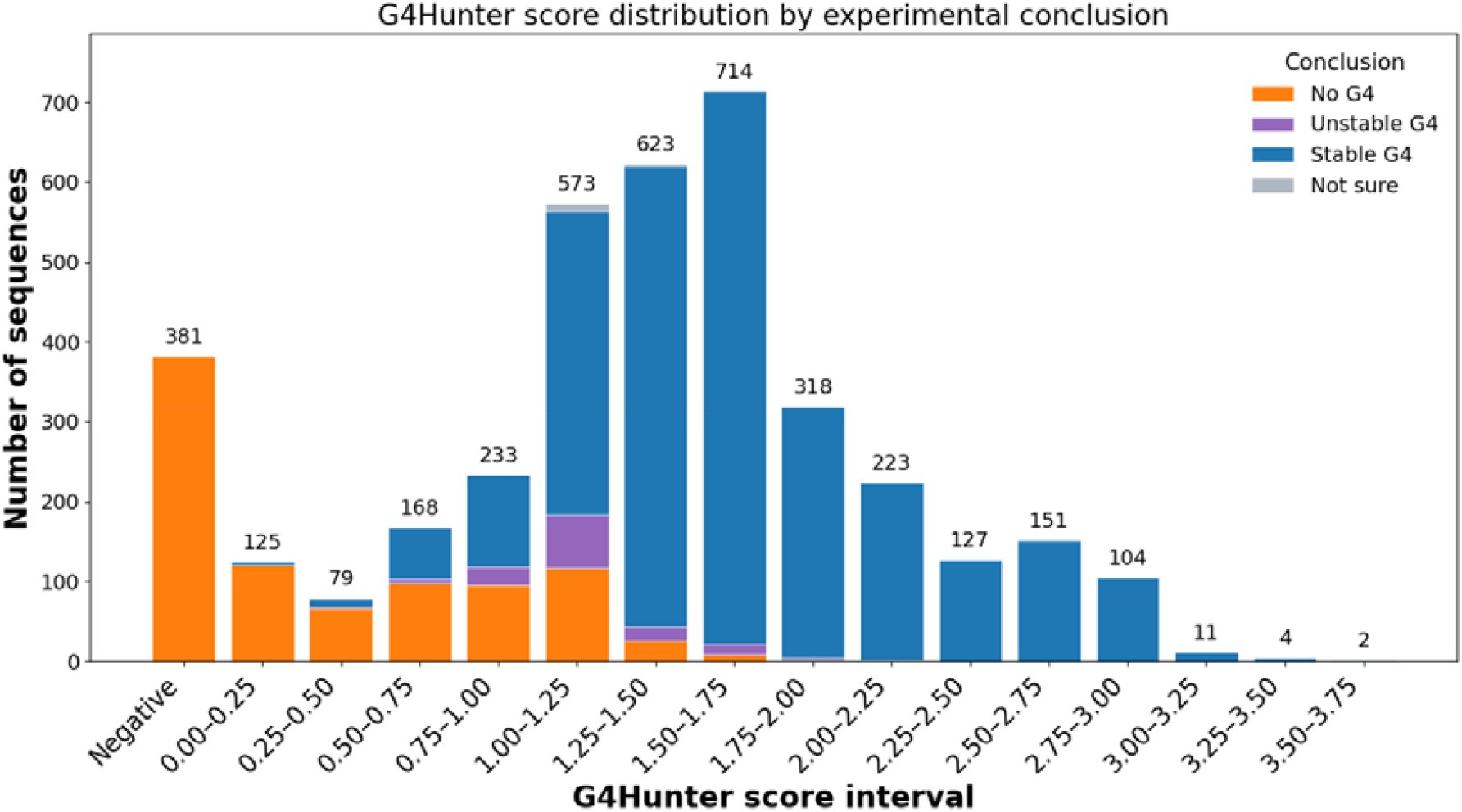
**Distribution of G4Hunter max scores and experimental outcome** (forming a stable G4, a G4 with a T_m_ below 37°C, or non-G4-forming in blue, purple and orange, respectively).

In addition, for the G4-forming sequences for which a T_m_ could be determined (*n*=1196; 799 of them currently validated used to generate **Figure 5**), a fair correlation (Spearman ρ= 0.6) could be found between T_m_ and G4Hunter score. G4Hunter is therefore not only an excellent predictor of G4 formation, but also a fair predictor of stability: the higher the score, the higher the chance that a G4 is actually formed and thermally stable.

**Figure 5:**
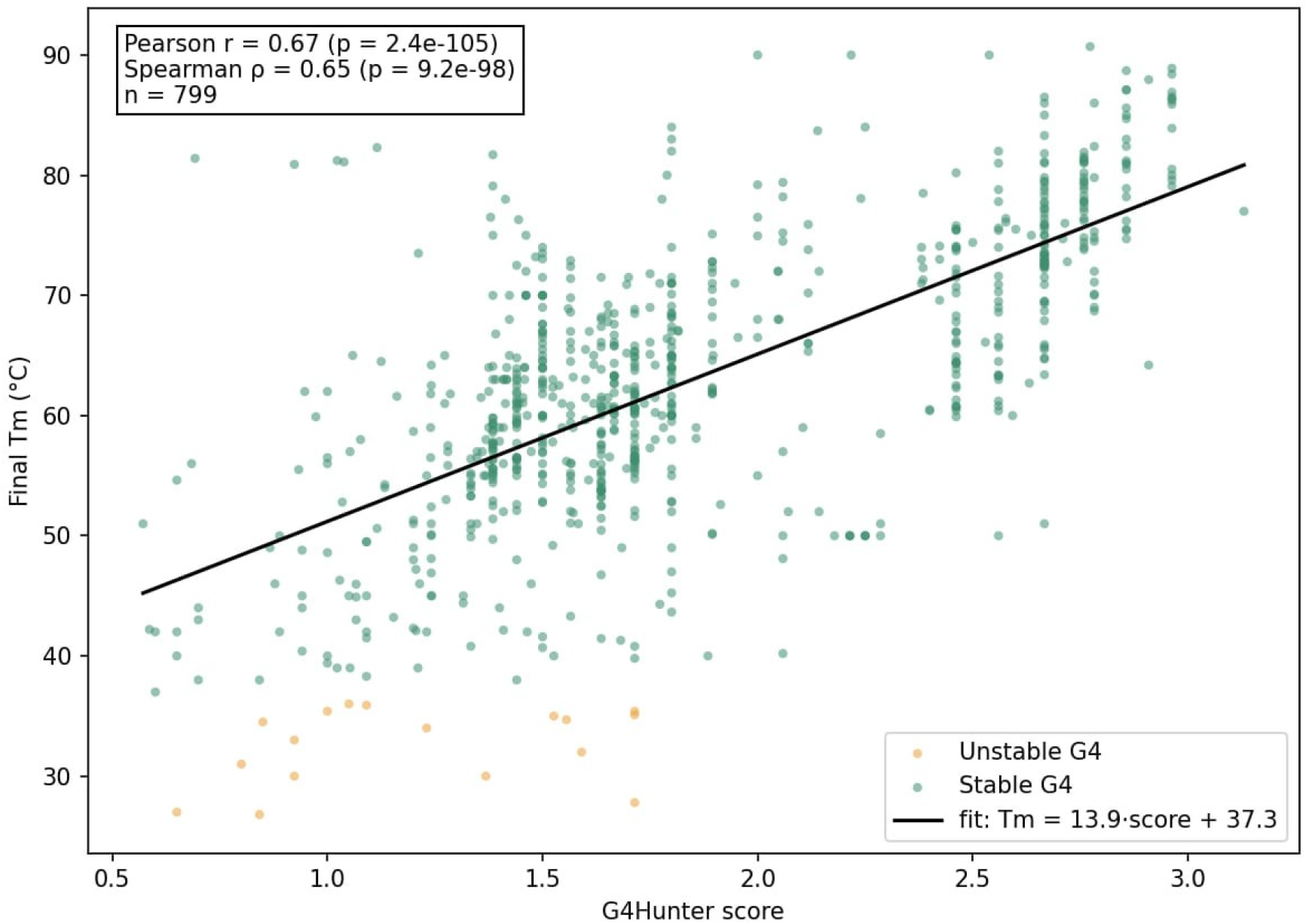
**T**_***m***_ **(in °C) versus** G4Hmax **(G4Hunter** “max” score) for a panel of 799 G4s.

## DISCUSSION

### Comparison with existing databases

Several databases have been developed to catalogue G-quadruplexes (see **Table 1)**.

G4All distinguishes itself by its focus on short, experimentally validated sequences paired with single-stranded or hairpin controls, a combination absent in existing resources. A number of databases are related to G-quadruplexes. G4LDB and G4IPDB report G4 ligands and G4-binding proteins, respectively. DSSR-G4DB lists G4 high resolution structures. G4RNA collects data for RNA sequences only. Databases such as QuadBase2 and G4IPDB primarily compile putative G4-forming sequences identified *in silico* across genomes, emphasizing predicted rather than experimentally confirmed structures. In contrast, G4All prioritizes laboratory-validated DNA and RNA sequences, providing a gold-standard dataset for benchmarking experiments and computational tools. Furthermore, unlike genome-wide repositories that catalogue G4 motifs in their native contexts, G4All focuses on short, synthetic sequences ideal for controlled *in vitro* assays, while its inclusion of non-G4 single-stranded controls enables rigorous discrimination between G4-specific and non-specific effects. This curated, dual-dataset approach positions G4All as a complementary resource that addresses critical gaps in reproducibility and experimental design.

### Advancing G4 research with standardized resources

The G4All database addresses a long-standing need in the nucleic acids community by providing a centralized, experimentally grounded repository of G4-forming DNA and RNA sequences, paired with rigorously selected non G4-forming controls. Unlike existing resources that primarily rely on *in silico* predictions or aggregate heterogeneous experimental data, G4All emphasizes curated, validated short (mostly synthetic) sequences with clear metadata on experimental conditions, structural topology, and thermal stability. We have chosen to retain data collected in potassium conditions (around 100 mM K^+^) at near-neutral pH (between 6.0 and 7.9). This standardization is critical for reproducible research, enabling direct comparisons across studies and reducing the variability that has historically complicated G4 characterization. By including both DNA and RNA sequences, G4All also bridges a gap between these two classes of nucleic acids, facilitating cross-disciplinary insights into G4 biology.

The inclusion of non G4-forming controls (single strands, hairpins, as well as other non-canonical nucleic acid structures such as i-DNA) is a defining feature of G4All. These sequences, which lack G4-forming ability under physiological conditions, provide essential benchmarks for experimental design, assay validation, and computational model training. This dual approach empowers researchers to distinguish G4-specific effects from non-specific interactions, a distinction that is particularly valuable in high-throughput screens and therapeutic target validation.

### Current limitations

The current composition of G4All reflects a notable bias toward DNA sequences, with significantly fewer RNA entries. This disparity mirrors historical trends in G4 research, where DNA G4s have been more extensively characterized due to their relative ease of DNA synthesis and manipulation, and the earlier development of DNA-specific detection methods. However, the biological and biomedical relevance of RNA G4s—particularly in post-transcriptional regulation, translation control, and viral genomes—is increasingly recognized. To address this imbalance, future versions of G4All will prioritize the curation of RNA G4 sequences, with a focus on those validated under physiological conditions and in cellular contexts, ensuring the database evolves to support the growing demand for RNA-focused applications in areas such as aptamer design, RNA-based therapeutics, and dynamic RNA structurome studies. For the DNA or RNA sequences with highly discordant Tm values reported under nearly identical conditions, additional experiments may be envisioned to solve this discrepancy.

The sequences curated in G4All reflect several sampling biases that influence their representativeness of *in vivo* G4 landscapes. A prominent example is the overrepresentation of sequences conforming to the consensus motif (G□N□□□G□N□□□G□N□□□G□) used in Quadparser (22), which has historically guided G4 prediction and experimental validation. However, this motif excludes many G4s with longer loops, bulges, mismatches, or variable loop compositions—features increasingly recognized in functional genomic and transcriptomic G4s, and can be predicted by G4Hunter (5,23). Nearly half the G4-forming stable G4s escape this Quadparser consensus, as well as the vast majority of unstable G4s (**Figure 6**). On the other hand, Quadparser generates very few false positives (sequences predicted to form a G4 but actually unable to do so).

**Figure 6:**
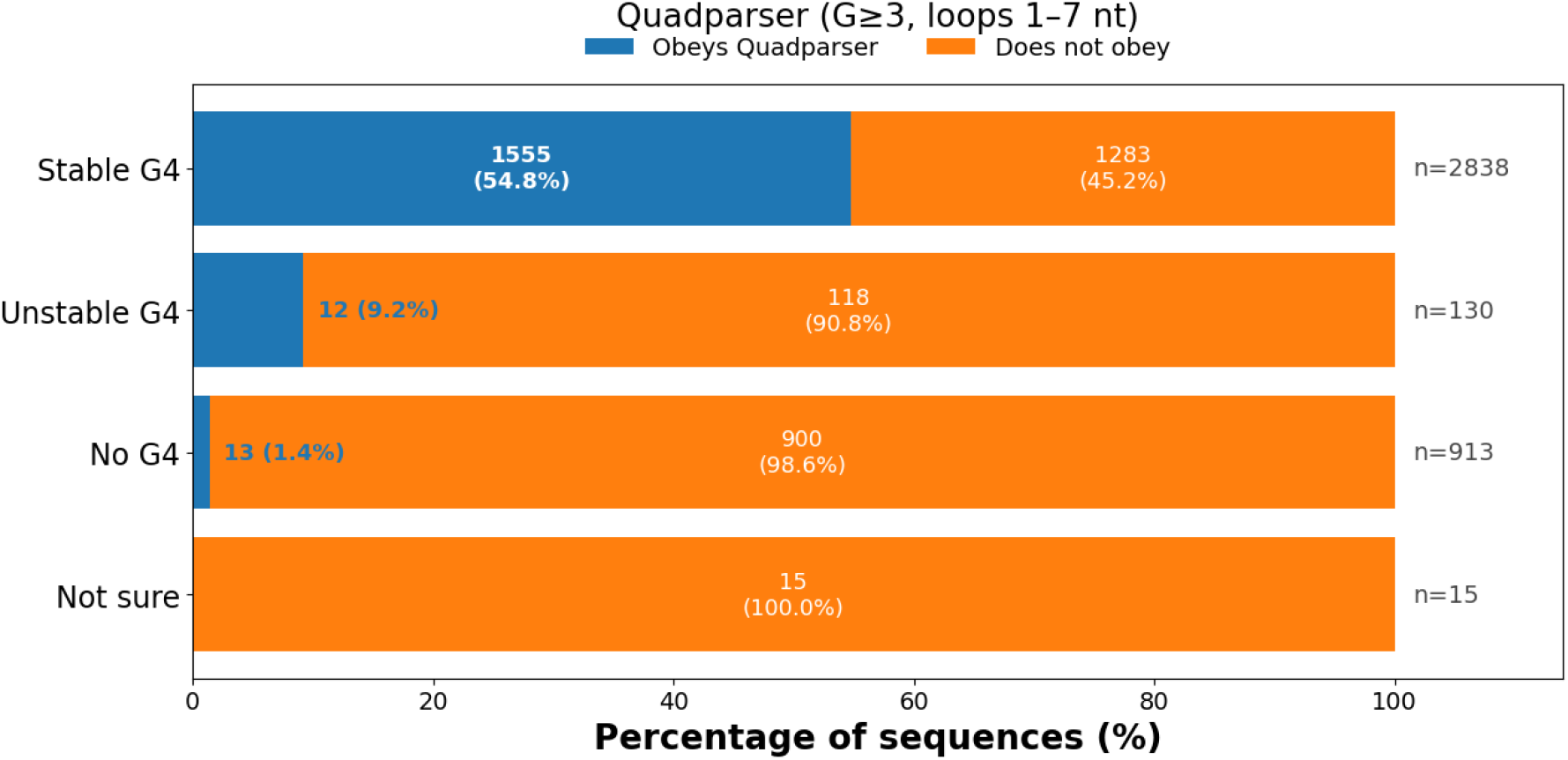
Quadparser sequences. In each category, the number of sequences obeying a classical Quadparser consensus (G□N□ □ □G□N□ □ □G□N□ □ □G□) is indicated, and the fraction shown in blue, while the fraction escaping it is shown in orange.

Additionally, the database may be biased toward highly stable sequences that are easier to characterize experimentally, as well as toward DNA over RNA and parallel topologies over antiparallel or hybrid forms, due to historical experimental preferences. Sequences from model organisms or specific genomic contexts (*e*.*g*., telomeres, promoter regions) may also be overrepresented. While these biases reflect the current state of the field, they underscore the need for future expansions of G4All to include diverse, non-canonical G4 structures validated under physiological conditions, ensuring broader applicability to real-world biological systems.

Another bias comes from the reluctance to publish negative results: there are probably more sequences for which G4 formation was tested but infirmed. This is unfortunate, as these negative controls are precious to train models able to predict G4 propensity. This bias is manifested by the relative paucity of sequences with a G4Hunter score between 0 and 1, and by the significantly larger number of G4-forming rather than non G4-forming sequences (three times more). G4 formation is unlikely, but not impossible, for sequences with a relatively low score (between 0.5 and 1). More motifs with scores in that range are going to be collected by our group, privileging natural rather than model sequences, in order to minimize this bias.

Another bias we identified is related to melting temperatures, which were found for “only” ≈ 1200 sequences among more than 2800 G4-forming motifs (41.83%). Many of the G4s reported here possibly exhibit a high thermal stability, making accurate T_m_ determination impossible. Yet, one could simply check if the structure remains folded at high temperature, for example by circular dichroism. In that case, we would attribute an arbitrary high T_m_ value (*e*.*g*., 95°C or more) to these sequences.

Another critical but often overlooked issue in G4 characterization is the molecularity of the observed structure. While most studies implicitly assume that a G4-forming sequence folds intramolecularly (within a single strand), this is rarely formally demonstrated. In reality, sequences can also form intermolecular G4s through the association of multiple strands, particularly at high concentrations or under specific ionic conditions. This distinction has profound implications: intramolecular G4s are more likely to form *in vivo* at physiological concentrations, whereas intermolecular G4s may either represent artifacts of *in vitro* conditions or reflect specific biological contexts where long-range contacts between G-runs allow G4 formation. To address this, G4All encourages contributors to provide experimental evidence for molecularity—such as concentration-dependence studies, crosslinking data, or single-molecule analyses—where available, and flags entries where this critical parameter remains undetermined.

While melting temperature (T_*m*_) provides a useful *in vitro* metric for G4 thermal stability, it is an imperfect proxy for physiological stability. T_*m*_ values are typically measured under simplified buffer conditions that do not recapitulate the complex cellular environment, where molecular crowding, heterogeneous ion compositions, and interactions with proteins or other biomolecules can either stabilize or destabilize G4s. Moreover, T_*m*_ reflects stability at equilibrium but fails to capture kinetic stability, dynamic folding–unfolding behaviour, or competition with alternative secondary structures that occur *in vivo*. In addition, T_*m*_ values could be extracted for only 41.83% of the G4 forming sequences analysed. The absence of T_*m*_ values for certain G4-forming sequences in G4All may reflect several factors: The original studies may not have measured or reported thermal stability, technical limitations may have prevented accurate determination (often because the G4 is very stable), or the melting process was complex or multiphasic. Thus, while T_*m*_ annotations in G4All offer valuable comparative insights, they should be interpreted alongside cellular validation data when available.

Unlike T_*m*_, which merely indicates the temperature at which 50% of the structure is denatured, the Gibbs free energy (ΔG) at physiological temperature (e.g., 37°C) directly quantifies the thermodynamic stability of G4 formation under biological conditions. ΔG integrates both enthalpic and entropic contributions to provide a temperature-specific measure of stability, reflecting the actual folded population in a cellular environment. This parameter is therefore more predictive of *in vivo* behaviour, enabling better estimation of G4 prevalence, kinetic accessibility, and competition with alternative structures or binding partners under native conditions. Unfortunately, reliable ΔG values have only been determined for a very small number of sequences.

### Future developments

To ensure G4All remains a dynamic and indispensable resource, we have outlined several key developments for the near future. This database will be constantly expanded by incorporating new experimental data: We will continuously integrate new sequences validated by biophysical or biochemical techniques, as well as user-submitted datasets following a peer-reviewed curation process. We will not only focus our efforts on human sequences, and the database actually includes motifs found in other organisms as well. A new public release will be made every trimester, and each version of the database will be labeled as “G4All_2026_Q3” for example, for the version released in August 2026. Future versions will incorporate additional topological classifications as well as thermodynamic parameters (*e*.*g*., ΔG°), and functional annotations (*e*.*g*., origin, associations with gene expression, disease mutations, or epigenetic marks). A submission portal will allow researchers to contribute new sequences, experimental protocols, and negative controls, fostering a collaborative ecosystem. Submitted data will undergo expert review to maintain high standards of quality and reproducibility. G4All will serve as a training and testing ground for predictive models of G4 formation. We plan to develop and host pre-trained classifiers to help users identify novel G4 candidates in custom sequences or genomes. Finally, we plan to experimentally re-assess G4 propensity for extreme outliers (G4-forming sequences with a very low G4Hunter score, or non-G4 forming motifs with a very high score). We identified a few dozen candidate motifs for reevaluation.

### Applications and broader impact

G4All is designed to catalyze advancements across multiple domains of nucleic acid research and biotechnology. By providing a reliable foundation of validated sequences, G4All will accelerate studies into the biophysical properties, cellular functions, and regulatory mechanisms of G4s. Researchers can leverage the database to design targeted experiments, such as FRET-based folding assays, CRISPR-based G4 editing, or live-cell imaging of G4 dynamics. G4All’s curated dataset can aid in the rational design of G4-stabilizing ligands and the identification of druggable G4 motifs in pathological contexts. The inclusion of controls will also improve the specificity of drug screening assays. G4All also holds significant potential for aptamer (24), biosensors, and nanodevices development, which are emerging as powerful tools in diagnostics and synthetic biology. The intrinsic stability and structural diversity of G4 motifs make them ideal scaffolds for engineering high-affinity aptamers targeting proteins, small molecules, or cells. By providing a comprehensive collection of validated G4-forming sequences, G4All can accelerate the rational design of G4-based aptamers for biosensing, targeted drug delivery, and diagnostic applications, while the inclusion of non-G4 controls helps optimize specificity and reduce off-target binding. G4All can serve as a design blueprint for engineering stable, functional G4 structures with tailored properties, such as pH-responsive or ion-selective folding.

Finally, this database will support the benchmarking and refinement of G4 prediction algorithms, addressing current limitations in accuracy and context-awareness. By providing both positive and negative datasets, G4All enables supervised learning approaches to improve *in silico* G4 validation in genomic and transcriptomic data.

### Challenges and outlook

While G4All represents a significant step forward, challenges remain in the field. Context-dependent G4 formation, influenced by ionic conditions, molecular crowding, and protein interactions, complicates the extrapolation of *in vitro* data to *in vivo* settings. Future iterations of G4All will incorporate cellular, species- and tissue-specific validation to better capture these nuances. Additionally, the dynamic nature of G4s, which can interconvert between topologies or coexist with other secondary structures (*e*.*g*., hairpins, i-motifs), necessitates integrative approaches that combine structural, biochemical, and computational perspectives.

Looking ahead, we envision G4All as a living hub for the G4 community, evolving alongside technological and scientific advancements. By fostering open data sharing, interdisciplinary collaboration, and standardized methodologies, G4All aims to propel the field toward a deeper, more actionable understanding of G4 biology, both for fundamental discoveries and translational applications.

## ACKNOWLEDGEMENTS

J.L.M. thanks all his past and present collaborators and co-workers for providing data on G4 formation and stability.

## AUTHOR CONTRIBUTIONS

Parth Rajput: Methodology, Formal analysis, Validation, Data curation, Website creation, Figure preparation, Writing—original draft. Anne Cucchiarini: Methodology, Validation, Figure preparation, Data curation, Writing—review & editing. Dmitrii Trubestkoy: Methodology, Writing—review & editing. Yun Chen: Formal analysis, Methodology, Writing— review & editing. Giacomo Ferrari: Formal analysis, Methodology, Writing—review & editing. Muafia Baigum: Formal analysis, Writing—review & editing. Laurent Lacroix: Formal analysis, Methodology, Validation, Writing— review & editing. Jean-Louis Mergny: Conceptualization, Formal analysis, Data curation, Methodology, Validation, Writing—original draft.

## SUPPLEMENTARY DATA

None.

## CONFLICT OF INTEREST

None declared.

## FUNDING

This work was supported by recurrent grants from Institut national de la Santé et de la recherche médicale (Inserm), Centre National de la Recherche Scientifique (CNRS) and Ecole Polytechnique. Y.C. is the recipient of a Chinese Scholarship Council PhD grant (202308310081). J.L.M. acknowledges support from Fondation de l’Ecole Polytechnique and Agence de l’Innovation de Défense (AID) via the Centre Interdisciplinaire d’Etudes pour la Défense et la Sécurité (CIEDS) [project2023–Pathogens]. Funding for open access charge: CIEDS.

## DATA AVAILABILITY

All the data collected is available for download using the following link: https://doi.org/10.5281/zenodo.21216943

## Notes

### Competing Interest Statement

The authors have declared no competing interest.

## REFERENCES

1. Spiegel J, Adhikari S, Balasubramanian S. The Structure and Function of DNA G-Quadruplexes. Trends Chem. 2020;2:123–136.

2. Varshney D, Spiegel J, Zyner K, Tannahill D, Balasubramanian S. The regulation and functions of DNA and RNA G-quadruplexes. Nat Rev Mol Cell Biol. 2020;21:459–474.

3. Huang K, Zhang S, Wang H, Qu Y, Lu Y, Li R, et al. Autonomous biomedical research with an artificial intelligence agent. Science, 2026; in press. 10.1126/science.adz4351.

4. Mergny JL, Phan AT, Lacroix L. Following G-quartet formation by UV spectroscopy. FEBS Lett. 1998;435:74–78.

5. Bedrat A, Lacroix L, Mergny JL Reevaluation of G4 propensity with G4Hunter. Nucleic Acids Res. 2016;44:1746–1759.

6. Li Q, Xiang JF, Yang QF, Sun HX, Guan AJ, Tang, Y.-L. G4LDB: a database for discovering and studying G-quadruplex ligands. Nucleic Acids Res. 2013; 41, D1115–D1123.

7. Yang Q, Wang XR, Wang YH, Wu X, Shi R, Wang Y, et al. G4LDB 3.0: a database for discovering and studying G-quadruplex and i-motif ligands. Nucleic Acids Res. 2025;53:D91–D98.

8. Mishra SK, Tawani A, Mishra A, Kumar A. G4IPDB: A database for G-quadruplex structure forming nucleic acid interacting proteins. Sci. Rep. 2016;6:38144.

9. Zok T, Kraszewska N, Miskiewicz J, Pielacinska P, Zurkowski M, Szachniuk M. ONQUADRO: a database of experimentally determined quadruplex structures. Nucleic Acids Res. 2022;50:D253–D258.

10. Yu H, Qi Y, Yang B, Yang X, Ding Y. G4Atlas: a comprehensive transcriptome-wide G-quadruplex database. Nucleic Acids Res. 2023; 51, D126–D134.

11. Bourdon S, Herviou P, Dumas L, Destefanis E, Zen A, Cammas A, Millevoi S, Dassi E. QUADRatlas: the RNA G-quadruplex and RG4-binding proteins database. Nucleic Acids Res. 2023;51:D240–D247.

12. Yadav VK, Abraham JK, Mani P, Kulshrestha R, Chowdhury S. QuadBase: genome-wide database of G4 DNA--occurrence and conservation in human, chimpanzee, mouse and rat promoters and 146 microbes. Nucleic Acids Res. 2008;36:D408–D413.

13. Dhapola P, Chowdhury S. QuadBase2: web server for multiplexed guanine quadruplex mining and visualization. Nucleic Acids Res. 2016;44:W110–W114.

14. Qian SH, Shi MW, Xiong YL, Zhang Y, Zhang ZH, Song XM, et al. EndoQuad: a comprehensive genome-wide experimentally validated endogenous G-quadruplex database. Nucleic Acids Res. 2024;52:D72–D80.

15. Kikin O, Zappala Z, D’Antonio L, Bagga PS. GRSDB2 and GRS_UTRdb: databases of quadruplex forming G-rich sequences in pre-mRNAs and mRNAs. Nucleic Acids Res. 2008;36:D141–D148.

16. Zhang R, Lin Y, Zhang CT. Greglist: a database listing potential G-quadruplex regulated genes. Nucleic Acids Res. 2008;36:D372–D376.

17. Garant JM, Luce MJ, Scott MS, Perreault JP. G4RNA: an RNA G-quadruplex database. Database, 2015;2015:bav059.

18. Vannutelli A, Schell LLN, Perreault JP, Ouangraoua A. GAIA: G-quadruplexes in alive creature database. Nucleic Acids Res. 2023;51:D135–D140.

19. Zhong H, Dong M, Gao F. G4Bank: A database of experimentally identified DNA G-quadruplex sequences. Interdiscip. Sci. Comput. Life Sci. 2023;15:363–373.

20. Cer RZ, Bruce KH, Mudunuri US, Yi M, Stephens RM. Non-B DB v2.0: a database of predicted non-B DNA-forming motifs and its associated tools. Nucleic Acids Res. 2013;41:D508–D513.

21. Chantzi N, Nayak A, Baltoumas FA, Aplakidou E, Liew SW, Galuh JE, et al. Quadrupia provides a comprehensive catalog of G-quadruplexes across genomes from the tree of life. Genome Res. 2025; 35, 2578–2600.

22. Kikin O, D’Antonio L, Bagga PS. QGRS Mapper: a web-based server for predicting G-quadruplexes in nucleotide sequences. Nucleic Acids Res. 2006; 34:W665–W669.

23. Brazda V, Kolomaznik J, Lysek J, Bartas M, Fojta M, Stastny J., et al. G4Hunter web application: a web server for G-quadruplex prediction. Bioinformatics 2019;35:2865–2867.

24. Cucchiarini A, Dobrovolná M, Brázda V, Mergny JL Analysis of quadruplex propensity of aptamer sequences. Nucleic Acids Res. 2025;53:gkaf424.

